# GigaSOM.jl: High-performance clustering and visualization of huge cytometry datasets

**DOI:** 10.1101/2020.08.03.234187

**Authors:** Miroslav Kratochvíl, Oliver Hunewald, Laurent Heirendt, Vasco Verissimo, Jiří Vondrášek, Venkata P. Satagopam, Reinhard Schneider, Christophe Trefois, Markus Ollert

## Abstract

**Background:** The amount of data generated in large clinical and phenotyping studies that use single-cell cytometry is constantly growing. Recent technological advances allow to easily generate data with hundreds of millions of single-cell data points with more than 40 parameters, originating from thousands of individual samples. The analysis of that amount of high-dimensional data becomes demanding in both hardware and software of high-performance computational resources. Current software tools often do not scale to the datasets of such size; users are thus forced to down-sample the data to bearable sizes, in turn losing accuracy and ability to detect many underlying complex phenomena.

**Results:** We present GigaSOM.jl, a fast and scalable implementation of clustering and dimensionality-reduction for flow and mass cytometry data. The implementation of GigaSOM.jl in the high-level and high-performance programming language Julia makes it accessible to the scientific community, and allows for efficient handling and processing of datasets with billions of data points using distributed computing infrastructures. We describe the design of GigaSOM.jl, measure its performance and horizontal scaling capability, and showcase the functionality on a large dataset from a recent study.

**Conclusions:** GigaSOM.jl facilitates utilization of the commonly available high-performance computing resources to process the largest available datasets within minutes, while producing results of the same quality as the current state-of-art software. Measurements indicate that the performance scales to much larger datasets. The example use on the data from an massive mouse phenotyping effort confirms the applicability of GigaSOM.jl to huge-scale studies.

**Key points:**

- GigaSOM.jl improves the applicability of FlowSOM-style single-cell cytometry data analysis by increasing the acceptable dataset size to billions of single cells.
- Significant speedup over current methods is achieved by distributed processing and utilization of efficient algorithms.
- GigaSOM.jl package includes support for fast visualization of multidimensional data.

## 1 Background

Advances in single-cell technologies, such as Mass Cytometry (CyTOF), Single-Cell RNA Sequencing (scRNA) and Spectral Flow Cytometry [1, 2, 3], provide deep and comprehensive insights into the complex mechanism of cellular systems, such as immune cells in blood, tumor cells and their microenvironments, and various microbiomes, including the single-celled marine life ecosystems. Mass cytometry and spectral cytometry have enabled staining of the cells with more than 40 different markers to discover cellular differences under multiple conditions. The samples collected in recent studies often contain millions of measured cells (events), resulting in large and high-dimensional datasets. Traditional analysis methods, based on manual observation and selection of the clusters in 2D scatter-plots, is becoming increasingly difficult to apply on data of such complexity: For high-dimensional data, this procedure is extremely laborious, and the results often carry researcher or analysis bias [4].

Various dimensionality reduction, clustering, classification and data mining methods have been employed to aid with the semi- or fully-automated processing, including the neural networks [5], various rule-based and tree-based classifiers in combination with clustering and visualization [6, 7], or locality-sensitive and density-based statistical approaches [8]. However, computational performance of the algorithms, necessary for scaling to larger datasets, is often neglected, and the available analysis software often relies on various simplifications (such as downscaling, which impairs the quality and precision of the result) required to process large datasets in reasonable time, without disproportionate hardware requirements.

To improve the performance, the underlying algorithm of FlowSOM [9] introduced a combination of the Self-Organizing-Maps (SOMs) by Kohonen [10] and metaclustering, which allowed efficient and accurate clustering of millions of cells [11]. FlowSOM is currently available as an R package that became an essential part of many workflows, analysis pipelines and software suites, including FlowJo and Cytobank® [12]. Despite of the advance, the amount of data generated in large research-oriented and clinical studies frequently grows to hundreds of millions of cells, processing of which requires not only the efficiency of the algorithm, but also a practical scalable implementation.

Here, we present GigaSOM.jl, an implementation of the SOM-based clustering and dimensionality-reduction functionality using the Julia programming language [13]. Compared to FlowSOM, GigaSOM.jl provides two major improvements: First, it utilizes the computational and memory resources efficiently, enabling it to process datasets of size larger than 10^8^ cells on commonly available hardware. Second, the implementation provides horizontal scaling support, and can thus utilize large high-performance computing clusters (HPC) to gain improvements in speed and tangible dataset size, allowing to process datasets with more than 10^10^ cells in the distributed environment. Additionally, the implementation in Julia is sufficiently high-level for allowing easy extensibility and cooperation with other tools in Julia ecosystem. Several technical limitations imposed by the R-wrapped implementation in the C programming language of FlowSOM are also overcome.

## 2 Methods

The Kohonen Self-Organizing-Map (SOM) algorithm [10] is a kind of simplified neural network with a single layer equipped with a topology. The task of the SOM training is to assign values to the neurons so that the training dataset is covered by neighborhoods of the neurons, and, at the same time, that the topology of the neurons is preserved in the trained network. A 2-dimensional grid is one of the most commonly used topologies, because it simplifies interpretation of the results as neuron values positioned in the 2-dimensional space, and related visualization purposes (e.g. EmbedSOM [14]). At the same time, the trained network can serve as a simple clustering of the input dataset, classifying each data point to its nearest neuron.

### 2.1 Batch SOM training

The original SOM training algorithm was introduced by Kohonen [15]. The map is organized as a collection of randomly initialized vectors (called *codebook*, with weights *W* (1)). The training proceeds in iterations (indexed by time *t*), where in each iteration a randomly selected data point in the dataset is used to produce an updated codebook as

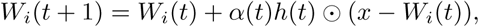

where *α* is the learning rate parameter, *i* is the neuron nearest to the randomly selected data point *x*, and *h* is the vector of topological distances of the codebook vectors to the best matching unit. The learning has been shown to converge after a predictable number of iterations if *α* and neighborhood size in *h* and topological neighborhood size are gradually lowered [10].

A more scalable variant of the algorithm can be obtained by running the single updates in batches where the values of *x* are taken from the whole dataset at once; which can be expressed in matrix form

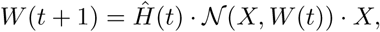

where 𝒩(*X, W* (*t*)) is a binary matrix that contains 1 at position *i, j* if and only if *W*_*i*_(*t*) is the closest codebook vector to *X*_*j*_, and *Ĥ* (*t*) is a distance matrix of the codebook in 2D map topology with rows scaled to sum 1. Notably, the algorithm converges in the same cases as the online version [16], and may be viewed as a generalized version of k-means clustering, which is obtained by setting *H*(*t*) = *I*.)

Implementations of the batch training may employ several assumptions that are not available with the online training:

- computation of 𝒩 can employ a pre-built spatial indexing structure on *W* (*t*), which is constant for the whole batch,
- all computations involving *X* can be sliced and parallelized (moreover, because the accesses to *X* are not randomized, the implementation is more cacheefficient and more suitable for SIMD- and GPU-based acceleration)
- multiplication by *Ĥ* (*t*) can be associatively postponed to work only with the small codebook matrix, saving more than 50% required computation volume when compared to online training with large neighborhoods.

### 2.2 Distributed implementation of GigaSOM.jl

The officially registered GigaSOM.jl package is a flexible, horizontally scalable, HPC-aware version of the batch SOM training written in the Julia programming language. The language choice has allowed a reasonably high-level description of the problem suitable for easy customization, while still supporting the efficient low-level operations necessary for fast data processing. GigaSOM.jl contains a library of functions for loading the data from Flow Cytometry Standard (FCS) files, distributing the data across a network to remote computation nodes present in the cluster, running the parallelized computation, and to exporting and visualizing the results. The overall design of the main implemented operations is outlined in Figure 1. Example Julia code that executes the distributed operations is provided in Supplementary Listing S1.

**Figure 1:**
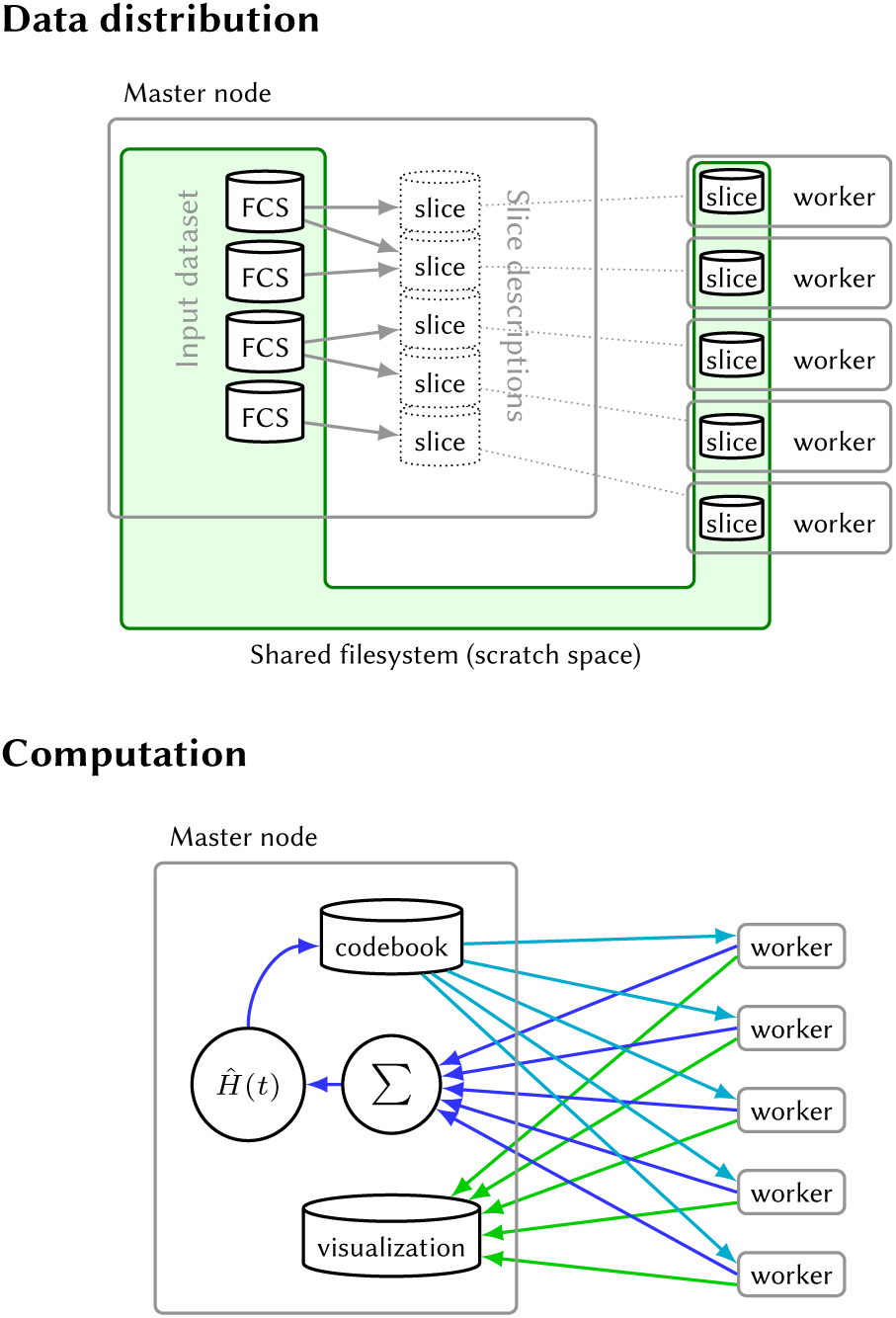
Architecture of GigaSOM.jl. Top: Data distribution process divides the available FCS files into balanced slices, individual workers retrieve their respective slice data using a shared storage. Below: The SOM learning and visualization processes require only a minimal amount of data transferred between the master and worker nodes; consisting of the relatively small codebook in case of SOM learning (blue arrows) and pre-rasterized graphics in case of visualization (green arrows).

#### 2.2.1 Data distribution procedure

Distributed computation process in GigaSOM is structured such as each computation node (‘worker’) keeps its own, persistent slice of the whole dataset, and the partial results from the nodes are aggregated by the master node. To establish this structure, GigaSOM implements a separate procedure that aggregates the input FCS files and creates a balanced set of slices equally distributed among the workers.

The distribution procedure is implemented as illustrated in Figure 1 (top): First, the master node reads the headers and sizes of individual FCS files, verifying their structure and determining the total number of stored data points. This is used to create minimal descriptions of dataset slices of equal size (each description consists only of 4 numbers of the first and last file and the first and last data point index), which are transferred to individual workers. Each worker interprets its assigned slice description, and extracts the part of the data from the relevant FCS files saved on a shared storage. The resulting slices may be easily exported to the storage and quickly imported again by individual workers, thus saving time if multiple analyses run on the same data (e.g., in case of several clustering and embedding runs with different parameters).

Importantly, a shared filesystem is usually one of the most efficient ways to perform data transfers in HPC environments, which makes the dataset loading process relatively fast. If a shared filesystem is not available, GigaSOM.jl also includes optional support for direct data distribution using the `Distributed.jl` package.

#### 2.2.2 Batch SOM implementation

After the nodes are equipped with the data slices, the batch SOM training proceeds as illustrated in Figure 1 (bottom):

1. The master node initializes the SOM codebook (usually by random sampling from available data).
2. The codebook is broadcast to all worker nodes. As the size of the usual codebook is at most several tens of kilobytes, data transfer speed does not represent a performance bottleneck in this case.
3. The workers calculate a partial codebook update on their data and send the results back to the master node.
4. Finally, the master node gathers the individual updates, multiplies the collected result by *Ĥ* (*t*), and continues with another iteration from step 2, if necessary.

Technically, the GigaSOM.jl implementation of steps 2–4 follows the structure of MapReduce data processing framework [17], which has allowed us to clearly separate the parallel processing implementation from actual computation primitives, and thus to improve the code maintainability. Apart from simplifying the implementation of various algorithm modifications, the MapReduce abstractions enable future transition to more complex data handling routines, such as the support for distributed parallel broadcast and reduction that is required for handling huge SOMs on very large number of workers (Collange et al. [18] provide a comprehensive discussion on that topic).

The time required to perform one iteration of the SOM training is mainly derived from the speed of the codebook transfer between nodes, and the amount of computation done by individual nodes. The current GigaSOM.jl implementation transfers all codebooks directly between the master node and the workers, giving time complexity 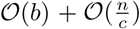 for *b* computation nodes equipped with *c* CPUs, working on a dataset of size *n*. This complexity can be improved to 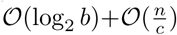 by using the aforementioned algorithms for parallel data broadcast and reduction, but we have not found a realistic dataset of size sufficient to gain any benefit from such optimization.

#### 2.2.3 Spatial indexing

Since the most computationally expensive step of the SOM training is the search for nearest codebook vectors for each dataset item (i.e., construction of the matrix 𝒩), we have evaluated the use of spatial indexing structures for accelerating this operation. GigaSOM.jl implementation can employ the structures available in the `NearestNeighbors` package, which include kd-trees and ball trees (also called vantage-point trees). [19, 20]

Although the efficiency of spatial indexing is vastly reduced with increasing dataset dimensionality, the measurements in section Results show that it can provide significant speedup with very large SOMs, even on data with more than 20 dimensions.

#### 2.2.4 Visualization support

To simplify visualization of the results, GigaSOM.jl includes a parallel reimplementation of the EmbedSOM algorithm in Julia [14], which quickly provides interpretable visualizations of the cell distribution within the datasets. EmbedSOM computes an embedding of the cells to 2-dimensional space, similarly as the popular t-SNE or UMAP algorithms [21, 22]. Unlike the usual dimensionality reduction algorithms, it uses the constructed SOM as a guiding manifold for positioning the individual points into the low-dimensional space, and achieves linear time complexity in the size of dataset. The parallel implementation of EmbedSOM is built upon the same distributed data framework as the batch SOMs — since EmbedSOM is designed to be trivially parallelizable, it can be run directly on the individual data slices, and gain the same speedup from parallel processing.

In order to aid the plotting of the EmbedSOM output, we have additionally implemented a custom scatterplot rasterizer in package `GigaScatter.jl`, which includes functions for quick plotting of large amounts of low-alpha points. To enable plotting of exceedingly large datasets, the rasterization can be executed in a distributed manner within the MapReduce framework, as shown in Supplementary Listing S1.

## 3 Results

The main result achieved by GigaSOM is the ability to quickly cluster and visualize datasets of previously unreachable size. In particular, we show that construction of a SOM from 10^9^ cells with 40 parameters can be performed in minutes, even on relatively small compute clusters with less than hundreds of CPU cores. The self-organizing map can be used to quickly dissect and analyze the samples, as with FlowSOM [**?**]. This performance achievement vastly simplifies the interactive work with large datasets, as the scientists can, for instance, try more combinations of hyperparameters and quickly get the feedback to improve the analysis and clustering of the data.

In this section, we first compare the output of GigaSOM.jl to that of FlowSOM, showing that the change in the SOM training algorithm has minimal impact on the quality of results. Further, we provide benchmark results that confirm that GigaSOM.jl scales horizontally, and details of the speedup achievable by employing spatial indexing data structures for acceleration of the nearest-neighbor queries. Finally, we demonstrate the achievable results by processing a gigascale dataset from a recent study by the International Mouse Phenotyping Consortium (IMPC) [23].

The presented performance benchmarks were executed on a Slurm-managed HPC cluster equipped with Intel®Xeon®E5-2650 CPUs; each node with 2 physical CPUs (total 24 cores) and 128GB of RAM. All benchmarks were executed several times, the times were measured as ‘real’ (wall-clock) time using the standard Julia timer facility. Measurements of the first runs were discarded to prevent the influence of caching and Julia just-in-time compilation; remaining results were reduced to medians.

### 3.1 Validation of clustering quality

To compare the GigaSOM.jl output with the one from Flow-SOM, we used a methodology similar to the one used by Weber and Robinson [11]. The datasets were first processed by the clustering algorithms to generate clusters, which were then assigned to ground truth populations so that the coverage of individual populations by clusters was reasonably high. The mean F1 score was then computed between the aggregated clusters and ground truth. Unlike Weber and Robinson [11], who use a complicated method of cluster assignment optimization to find the assignment that produces the best possible mean F1 score, we employed a simpler (and arguably more realistic) greedy algorithm that assigns each generated cluster to a population with the greatest part covered by that cluster.

The benchmark did not consider FlowSOM metaclustering [9], since the comparison mainly aimed to detect the differences caused by the modifications in SOM training.

For the comparison, we reused the datasets `Levine_13dim` and `Levine32_32dim` from the clustering benchmark [11]. In a typical outcome, most populations were matched by GigaSOM.jl just as well as by FlowSOM, as displayed in Figure 2 (detailed view is available in supplementary figure S1). Both methods consistently achieved mean F1 scores in the range of 0.65–0.7 on the `Levine_13dim` dataset and 0.81–0.84 on the `Levine_32dim` dataset for a wide range of reasonable parameter settings. In the tests, neither algorithm showed a significantly better resulting mean F1 score.

**Figure 2:**
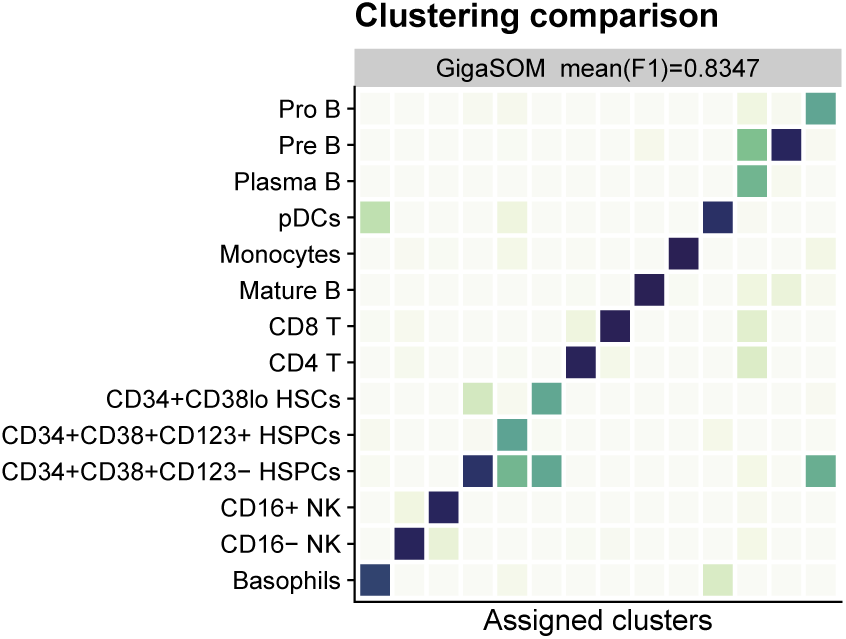
Comparison of GigaSOM.jl results with manual gating of the *Levine32* dataset. The confusion matrix is normalized in rows, showing the ratio of cells in each aggregate of GigaSOM-originating clusters that matches the cell types from manual analysis. Darker color represents better match. The mean F1 score is comparable to FlowSOM. A more comprehensive comparison is available in Supplementary Figure S1.

### 3.2 Scalable performance on large computer clusters

The benchmark of implementation scalability was performed as follows: A randomly generated dataset was distributed among the available computation nodes (workers) so that all CPUs are assigned an equal amount of data. For the benchmark, node counts as powers of two up to 256 have been chosen while the numbers of dataset parameters were chosen from multiples of 10 up to 50. The size of the dataset slice for a single node varied between 100, 200 and 300 thousand cells to verify the impact of data density in cluster. The dataset was then processed by the SOM training algorithm for SOM sizes 10×10, 20×20 and 40×40. The resulting SOMs were used for classifying the dataset into clusters (each input data point was assigned to a cluster defined by the nearest neighbor). An embedded view of the data was produced with the Julia implementation of EmbedSOM. All algorithms were also tested in variants where the naive search for nearest neighbors (or *k*-neighborhoods in case of EmbedSOM) was replaced by utilization of a spatial-indexing data structure, in particular by the kd-trees and ball-trees.

The scalability results are summarized in Figure 3: All three implemented algorithms scale almost linearly with the dataset size, the size of the SOM, and the dimension of the dataset. They reach an almost linear speedup with added compute capacity. In the case of SOM training, the required communication among the nodes caused only a negligible overhead; the majority of the computation pauses was caused by the random variance in execution time of computation steps on the nodes. The parallelized classification and embedding algorithms did not suffer from any communication overhead. Detailed benchmark results that show precise energy requirements of the training per processed data point, useful for deployment in large environments, are available in supplementary figure S2.

**Figure 3:**
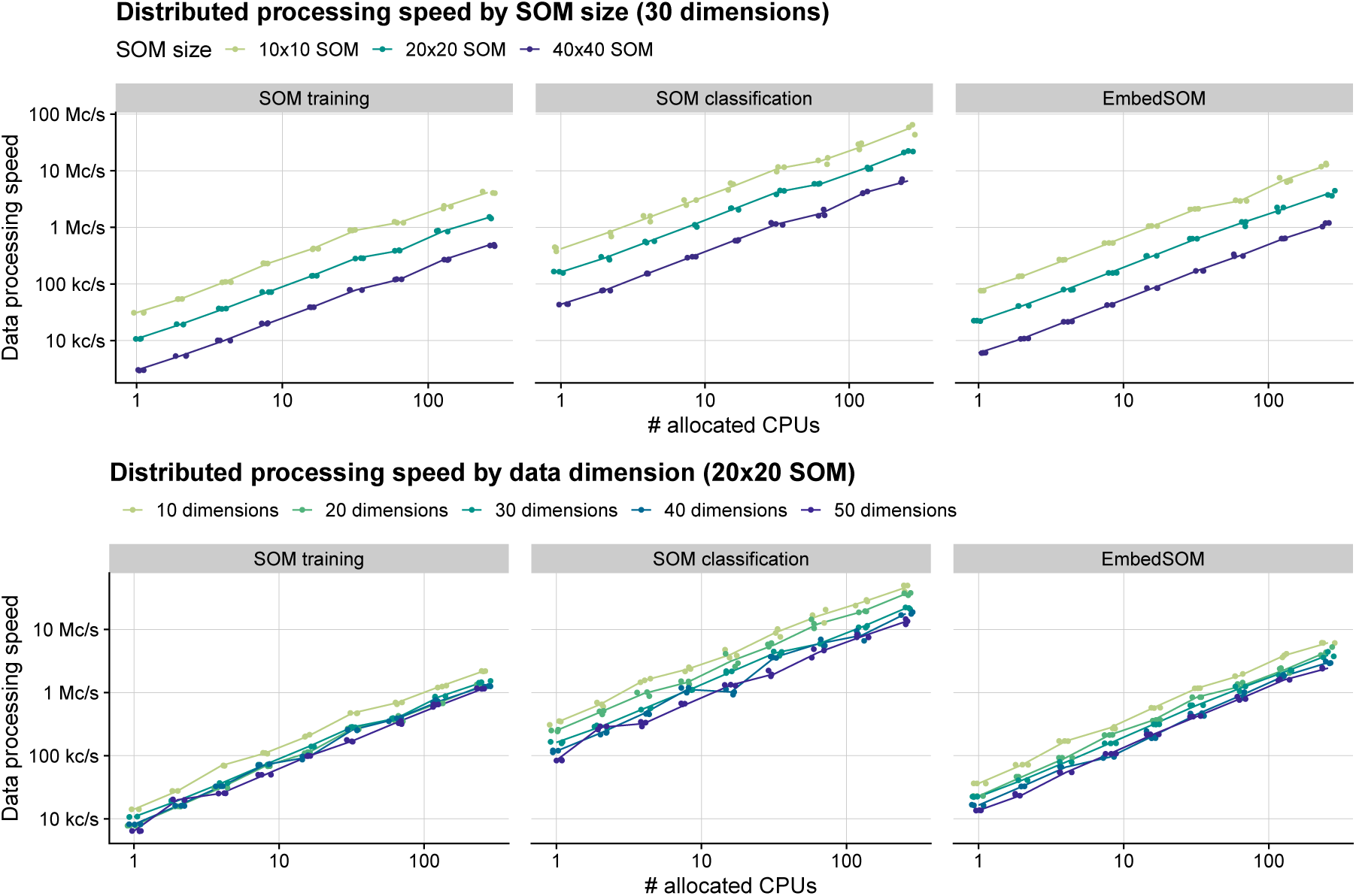
Performance dependency of distributed algorithms in GigaSOM on data dimensionality, SOM size and number of available workers. Data processing performance is displayed as normalized to median speed in cells per second (c/s).

Influence of the spatial indexing on the speed of various operations was collected as relative speedups (or slow-downs) when compared to a naive search. The results are displayed in Figure 4. We have observed that both kd-trees and ball-trees were able to accelerate some operations by a factor above 2×, but the use of spatial indexing suffered from many trade-offs that often caused performance decrease.

**Figure 4:**
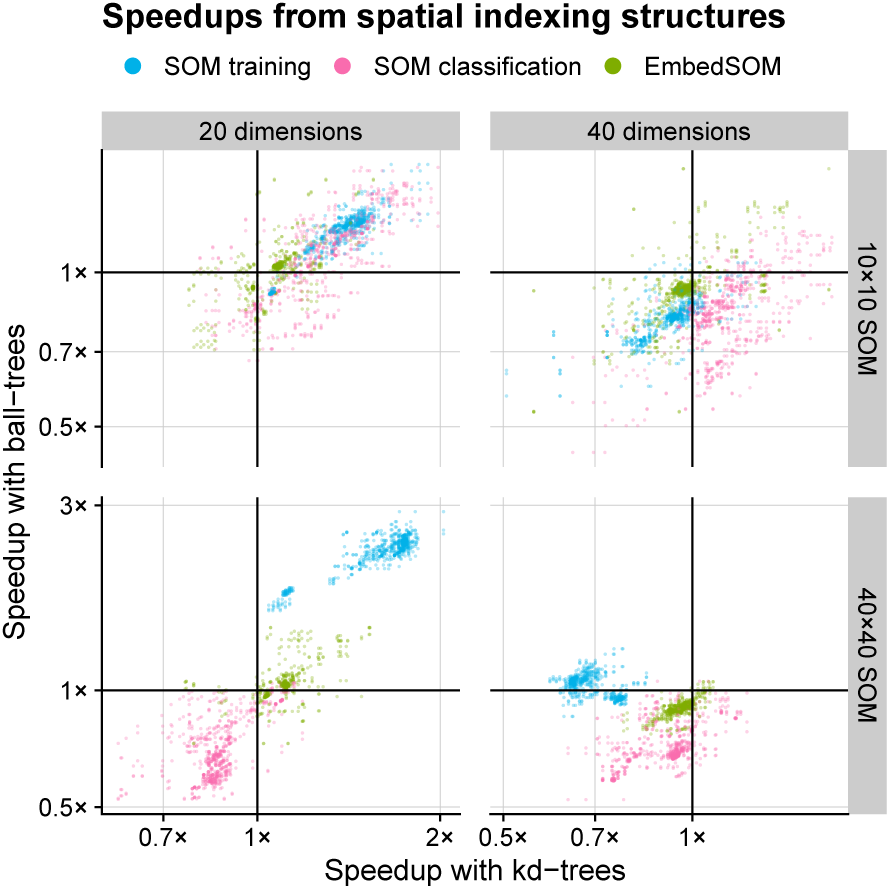
Effect of data indexing structures on GigaSOM performance. The plotted points show relative speedup of the algorithms utilizing kd-tress (horizontal axis) and ball-trees (vertical axis) compared to the brute-force neighbor search. Baseline (1× speedup) is highlighted by thick grid lines — a point plotted in the upper right quadrant represents a benchmark measurement that showed speedup for both kd-trees and ball-trees, upper left quadrant contains benchmark results where ball-trees provided speedup and kd-trees slowed the computation down, etc.

Most importantly, the cost of building the index has often surpassed the total cost of neighborhood lookups by the naive method, which is most easily observable on the measurements of ball-tree performance with smaller SOM sizes. Both trees have struggled to provide sufficient speedup in presence of higher dimensionality overhead (over 30), and had only negligible impact on the execution time of EmbedSOM computation, which was dominated by other operations.

On the other hand, it was easily possible to gain speedups around 1.5×for SOM training in most tests with lower dimension and large SOM, reaching 2.7×for a 20-dimensional dataset (typical for current flow cytometry) processed with large 40×40 SOM. From the results, it seems appropriate to employ the spatial indexing when the cost of other operations outweighs the cost of building the index, and the dimensionality overhead does not impede the efficiency of indexed lookup; in particular when training large SOMs of dimensionality less than around 30, and when data occupancy per node is sufficiently high. Detailed measurements for all SOM sizes and dataset dimensions are available in Supplementary Figure S3.

### 3.3 HPC analysis of previously unreachable dataset sizes

To showcase the GigaSOM.jl functionality on a realistic dataset, we have used a large dataset from the IMPC phenotyping effort [23] that contains measurements of mouse spleens by a standardized T-cell targeting panel. with individual cohorts containing genetically modified animals (typically a single-gene knockouts) and controls; total 2905 samples contain 1,167,129,317 individual cells. (The dataset is available from FlowRepository under the accession ID `FR-FCM-ZYX9`.)

The dataset was intentionally prepared by a very simple process — cell expressions were compensated, fluorescent marker expressions were transformed by the common *asinh* transformation with co-factor 500, and all dataset columns were scaled to *µ* = 0 and *σ* = 1. The resulting data were used to train a 32×32 SOM, which was in turn used to produce the embedding of the dataset (with EmbedSOM parameter *k* = 16), which was rasterized. The final result can be observed in Figure 5. The detailed workflow is shown in Supplementary Listing S1.

**Figure 5:**
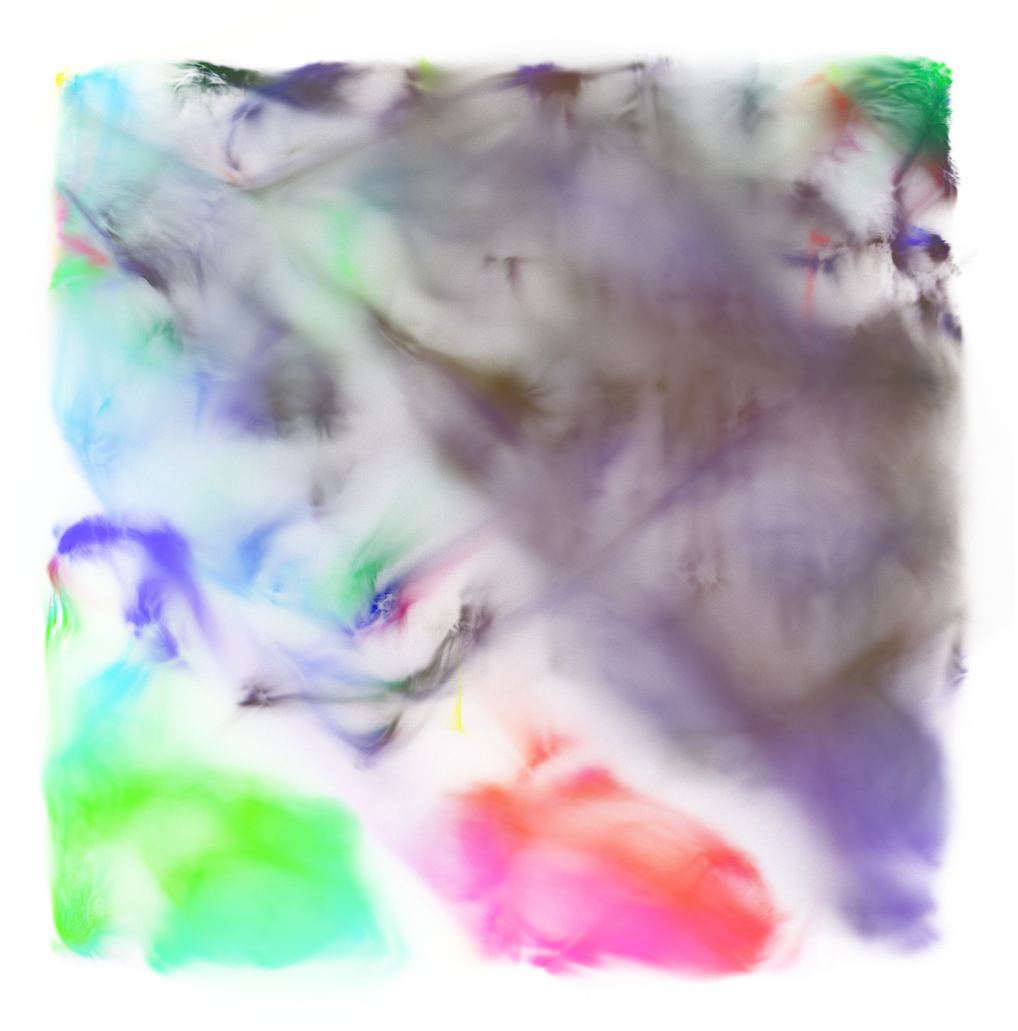
Raw IMPC Spleen T-cell dataset, processed by GigaSOM.jl and embedded by the Julia implementation of EmbedSOM. The figure shows an aggregate of 1,167,129,317 individual cells. Expression of three main markers is displayed in combination as mixed colors; CD8 in red, CD4 in green, and CD161 in blue. A more detailed, annotated version of the visualization is available in Supplementary Figure S4.

Notably, on a relatively small 256-core computer cluster (total 11 server nodes within a larger cluster managed by Slurm), the whole operation, consisting of Julia initialization, data loading (82.6GB of FCS files), SOM training for 30 epochs, embedding and export of embedded data (17.4GB) took slightly less than 25 minutes, and consumed at most 3GB of RAM per core. From that, each epoch of the parallelized SOM training took around 25 seconds, and the computation of EmbedSOM visualization took 3 minutes. Distributed plotting of the result was done using the `GigaScatter.jl` package; the parallel rasterization and combination of partial rasters took slightly over 4 minutes.

## 4 Conclusions

In this paper, we presented the functionality of GigaSOM.jl, a new, highly scalable toolkit for analyzing cytometry data with algorithms derived from self-organizing maps. The results conclusively show that GigaSOM.jl will support the growing demand for processing of huge datasets, and bolster the utilization of the HPC hardware resources that are becoming widely available for labs and universities.

The ability to process a gigascale dataset to a comprehensible embedding and precise, easily scrutinizable statistics in mere minutes may play a crucial role in both design and analysis methods of future cytometry experiments. We believe that the accessible and flexible nature of the GigaSOM.jl implementation in Julia programming language will also drive a transformation of other tools in the ecosystem towards the support of big data processing paradigms.

The resulting software is publicly available as a Julia package. The interoperability with the Julia ecosystem allows GigaSOM.jl to benefit from many other available scientific computing packages, which simplifies its deployment not only in cytometry, but also in other areas of research that employ self-organizing maps to extract information from large datasets.

## Data and software availability

All data and software is available under https://doi.org/10.17881/lcsb.z5vy-fa75.

- Package name: GigaSOM.jl
- Package home page: https://git.io/GigaSOM.jl
- Operating system(s): Portable to all Julia-supported platforms
- Programming language: Julia
- Other requirements: –
- License: Apache License v2.0

## Declarations

### Funding

MK and JV were supported by ELIXIR CZ LM2018131 (MEYS).

This work was supported by the Luxembourg National Research Fund (FNR) through the FNR AFR-RIKEN bilateral program (TregBar 2015/11228353) to MO, and the FNR PRIDE Doctoral Training Unit program (PRIDE/11012546/NEXTIMMUNE) to VV, RS and MO.

The Responsible and Reproducible Research (R3) team of the Luxembourg Centre for Systems Biomedicine is acknowledged for supporting the project and promoting reproducible research.

The experiments presented in this paper were carried out using the HPC facilities of the University of Luxembourg [24] (see https://hpc.uni.lu).

The project was supported by Staff Exchange programme of ELIXIR, the European life-sciences infrastructure.

### Author’s Contributions

Conceptualization: OH, LH, CT. Formal analysis, investigation, methodology: OH, MK, LH. Software: OH, MK, LH, VV. Funding acquisition, supervision: JV, VPS, RS, CT, MO. Validation: OH, MK. Visualization: MK. Writing: OH, MK. All authors participated in reviewing, editing and finalization of the manuscript.

