## Supplementary figures and code for "GigaSOM.jl: High-performance clustering and visualization of huge cytometry datasets"

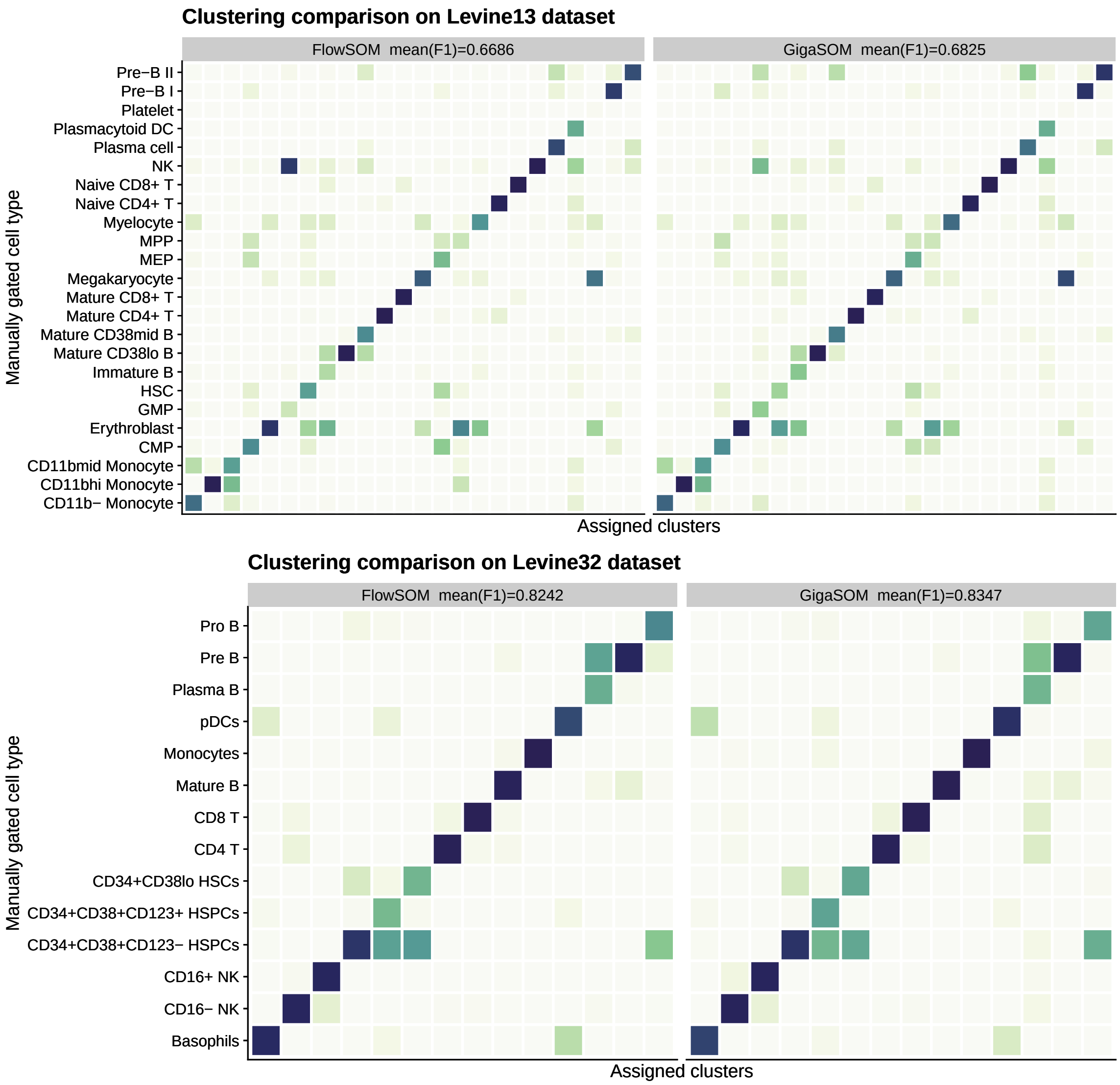


Figure S1: Results of clustering precision comparison with FlowSOM, on Levine13 and Levine32 datasets.

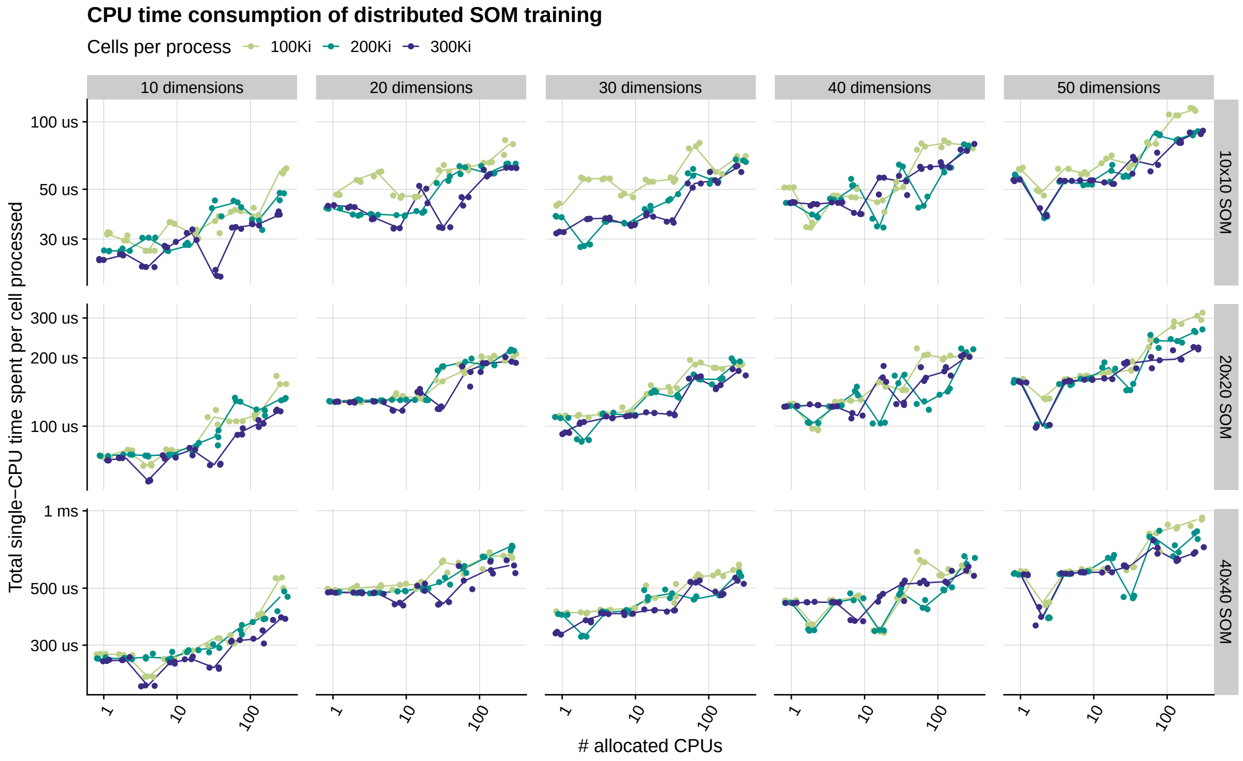


Figure S2: Detailed overview of the effect of horizontal scaling on the performance of SOM training in GigaSOM.jl. The color distinguishes different benchmarked levels of CPU occupancy (100 to 300 thousand cells loaded in memory per CPU).

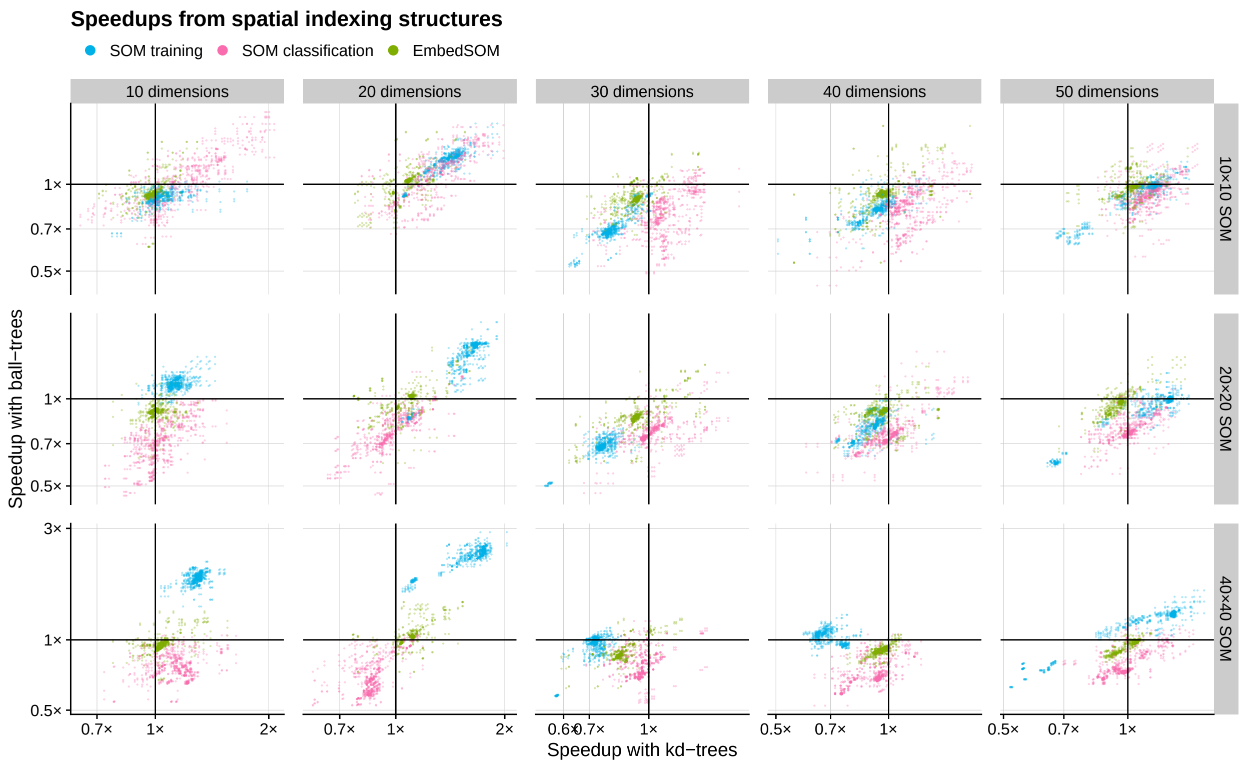


Figure S3: Detailed overview of the speedups provided by using spatial indexes for accelerating the nearest neighbor queries in implemented algorithms.

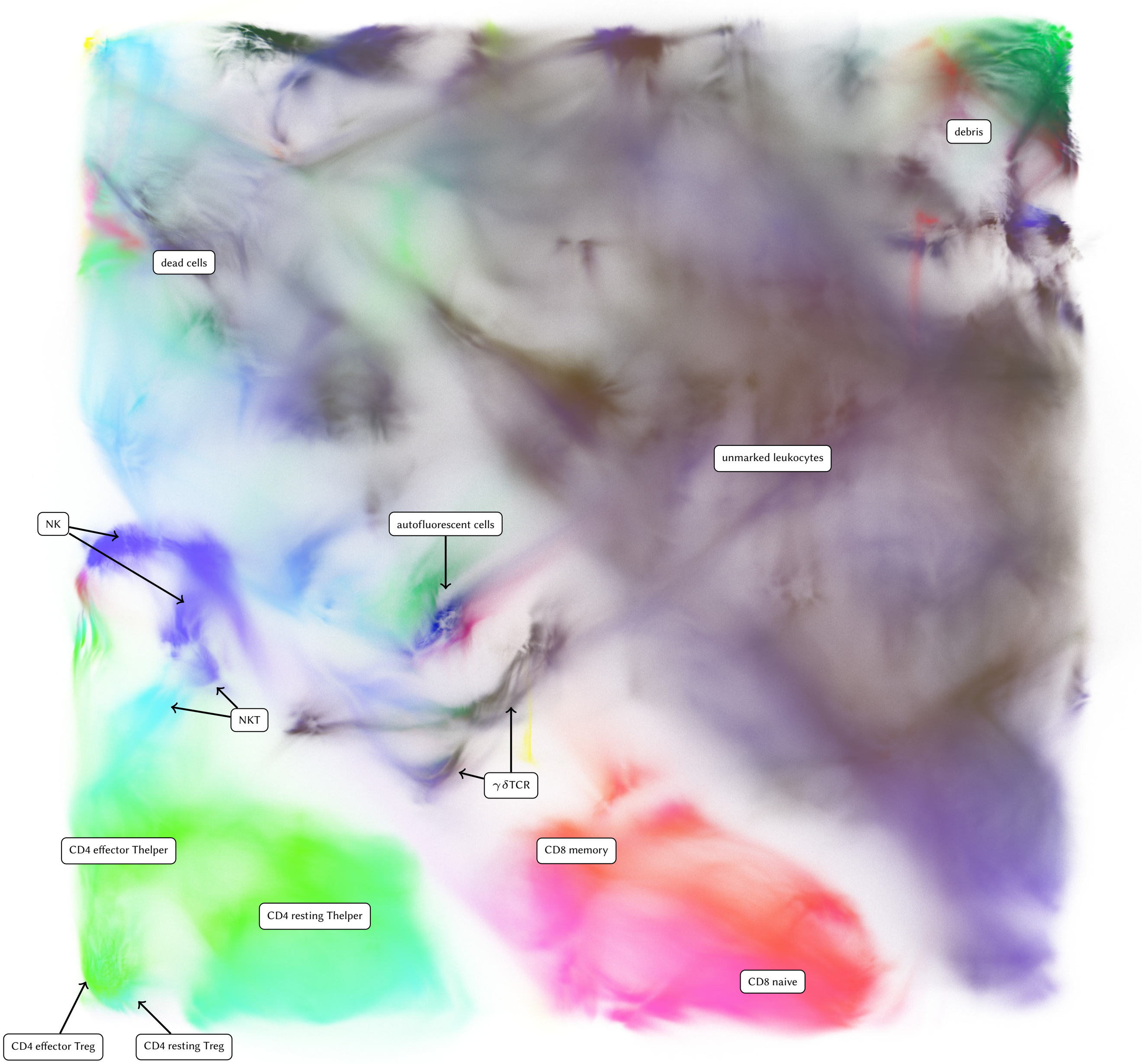


Figure S4: Larger version of Figure 5 — an embedding of more than 1 billion cells from the IMPC dataset (`FR-FCM-ZYX9`). The main lineage markers are highlighted as combinations of RGB colors: CD8 in red, CD4 in green and CD161 in blue. Annotations are made based on presence of several markers that are not highlighted in the embedding, mainly the Live/Dead marker, $\gamma\delta$TCR, CD25, GITR, and fluorescence scatter intensities.

```julia
using Distributed, GigaSOM, GigaScatter, ClusterManagers
import Glob, NearestNeighbors, Distributions

# find the file names to process
files = Glob.glob("TC_SPL_*.fcs")

# read the number of available workers from environment and start the worker processes
n_workers = parse(Int, ENV["SLURM_NTASKS"])
addprocs_slurm(n_workers, topology=:master_worker)

# load the required packages on all workers
@everywhere using GigaSOM
@everywhere using GigaScatter

# load the data to workers
dataset = loadFCSSet(:IMPCdata, files)

# collect the total number of cells from all workers
dataset_size = distributed_mapreduce(dataset, d->size(d,1), +)
@info "Loaded total $dataset_size cells."

# preprocess the distributed data
dselect(dataset, Vector(1:18)) # columns 1-18 contain interesting information
dtransform_asinh(dataset, Vector(7:18), 500.0) # transform the marker expressions
dscale(dataset, Vector(1:18)) # scale all columns

# train the SOM and run the embedding (this uses all available Slurm workers)
som = initGigaSOM(dataset, 32, 32)
som = trainGigaSOM(som, dataset, epochs=30,
                   rFinal=0.1, radiusFun=expRadius(-20.0),
                   knnTreeFun=NearestNeighbors.BallTree)
e = embedGigaSOM(som, dataset, k=16, output=:embeddedIMPC)

# setup constants for rasterization of the embedding
rasterSize = (2048, 2048)
alpha = 0.003
dist = Distributions.Normal()

# run the distributed rasterization
compound_raster = mixedRaster(distributed_mapreduce(
    [dataset, e],
    (d, e) -> begin
        # convert the normalized expressions to numbers between 0--1
        colors = hcat(
            Distributions.cdf.(dist, d[:,10]), # R: CD8 column
            Distributions.cdf.(dist, d[:,16]), # G: CD4 column
            Distributions.cdf.(dist, d[:,15]), # B: CD161 column
            fill(alpha, size(d,1)))
        # rasterize the local part of the data
        mixableRaster(rasterize(
            rasterSize,
            Matrix{Float64}(e'),
            Matrix{Float64}(colors'),
            xlim=(-2,33), ylim=(-2,33)))
    end,
    mixRasters))

# save the result to a PNG
savePNG("impc-embedding.png", compound_raster)
```

Listing S1: Complete code for Julia workflow that produces the embedding of the IMPC dataset (Figures 5 and S4).
